# Effect of a topically applied gel containing antibodies from bovine milk on the relative abundance of Staphylococci in the canine skin microbiome

**DOI:** 10.1101/2023.07.29.551104

**Authors:** Eline S. Klaassens, Silvia Auxilia, Mirna L. Baak, Radhika Bongoni, Catharina M.M. van Damme, Sannah Bijker, Nardy Dietz, Esther Winter, Femke Broere, Beatrix Foerster

## Abstract

Canine pyoderma is characterised by a predominant colonisation of the skin with *Staphylococcus pseudintermedius* and inflammatory symptoms associated with it. Immunoglobulins isolated from cow^’
s^s milk are neutralizing virulence factors and toxins of *S. pseudintermedius*. In a pilot study with four dogs with pyoderma we were able to show that the application of immunoglobulin gel leads to a decrease in the staphylococcal population and an increase in the diversity of the skin microbiome.

Due to the limited number of animals, the measurable effect is not statistically significant. Therefore, we are planning a follow-up study that includes more animals.

## 1. Introduction

The skin of dogs is inhabited by a rich and diverse microbiome. It shows a high variability between different individuals as well as different parts of the body. The skin microbiome plays an important role in skin health, but also in the pathophysiology and in the prevention of disease (Rodrigues Hoffmann et al., 2014).

Superficial pyoderma is a common medical condition affecting dogs, consisting of skin infection and inflammation. (Hillier et al., 2014; Loeffler and Lloyd, 2018). The primary pathogen in canine pyoderma is the gram positive, catalase-positive and facultative anaerobic bacterium *Staphylococcus pseudintermedius (S. pseudintermedius*), which is also a significant member of the normal flora in canines (Biberstein et al., 1984). In pyoderma samples, Staphylococcus spp. abundance has been shown to be increased, whereas *S. pseudintermedius* was the primary staphylococcal species found (Biberstein et al., 1984).

The ideal topical treatment for active pyoderma and to prevent recurrence of the disease should repress pathogenic Staphylococci without negatively influencing commensal skin bacteria. Therefore, preference should be given to narrow-spectrum topical antimicrobial agents wherever possible.

Topical antibiotics and antiseptics are often employed to inhibit colonization by pathogenic bacteria on the skin. In response to antibiotics, cutaneous bacterial populations exhibited an immediate and long-term shift in bacterial residents. These alterations can have critical implications for cutaneous host defence in the context of recolonization and a higher susceptibility to pathogenic staphylococci. Topical antiseptic treatments do not significantly alter skin bacterial community structure (SanMiguel et al., 2017) and even they have even been shown to increase the diversity of bacterial and fungal compositions (Chermprapai et al., 2019).

Bacteriophages as well as antimicrobial peptides are therapeutic options currently in development and specifically targeting single bacterial strains with little impact on the normal microbiome (Squires, 2021).

A bovine polyclonal antibody isolated from milk are binding virulence factors of Staphylococci. Our hypothesis was that this will diminish the pathogenicity of the bacteria and reduce their survival advantage in comparison to commensal bacteria. In addition, neutralising the virulence factors that damage skin and its immune system should result in more resilient skin that better fights off pathogenic bacteria. As a result, we expect a decrease in pathogenic bacteria and a more balanced skin microbiome.

In order to confirm the hypothesis, we investigated the influence of a bovine polyclonal antibody isolated from milk with activity against Staphylococcus on skin microbiota on lesioned canine skin as well as healthy skin compared to a placebo.

## 2. Methods

### Dog patients

Four dog patients with diffuse to generalised superficial pyoderma of equal distribution on both sides of the body were presented to the dermatology referral service of the clinic for companion animals of Utrecht University. The diagnosis of superficial pyoderma was based on clinical (Bensignor et al., 2016) and bacteriological findings and confirmed by cytology performed by the dermatology service (Budach and Mueller, 2012). *S. pseudintermedius* had to be present in the skin lesions as main pathogen, and this was confirmed by a bacterial culture and sensitivity test (sent to VMDC laboratory, Utrecht University). Dogs receiving systemic antimicrobials for the previous two weeks were not eligible. Additional topical treatments during the experiment were not permitted.

All dogs were privately owned. All experimental procedures were approved by the Utrecht University Animal Ethic committee as required under Dutch legislation and informed written consents were obtained prior from dog owners.

The study was conducted in a double blinded manner. The owners received two bottles of gel from the Utrecht University, Faculty of Veterinary Medicine, pharmacy department. One of them contained 1.5% (w/v) sodium carboxymethylcellulose containing 10 mg/ml immunoglobulin isolated from bovine milk as verum (with known activity against virulence factors of *S. pseudintermedius)* (Geh et al., 2019) while the other bottle was a placebo gel (1.5% (w/v) sodium carboxymethylcellulose). Both gels were applied twice a day for 14 sequential days, the placebo gel on one side and the immunoglobulin gel on the other affected side of each dog. The dogs were washed on day 0 before the start of the treatment with Malaseb® shampoo (Dechra Veterinary Products A/S, Uldum, Denmark), containing 2% miconazolenitrate and 2% chlorhexidinedigluconate.

### Microbiota skin assessment

On day 0 and 14, three skin samples were taken from different locations: a healthy area, an affected area on one side and an affected area on the other side of the dog. The samples were collected by sterile swabbing, using Epicentre Catch-AllTM swabs. The sterile swab was immersed into a tube with sterile 0.9 % NaCl-solution for moistening. Thereafter, the moistened swab was swept over the according skin area with constant and adequate pressure while rotating the swab for approximately 30 seconds. Subsequently, the swab head was placed into a collection tube cryovials pre-filled with 1 ml DNA stabilization buffer. The samples were stored in a freezer and within three months sent refrigerated to BaseClear B.V. in Leiden, the Netherlands, for DNA extraction.

### 16S and ITS amplicon sequencing of the skin microbiome

Paired-end sequence reads were generated using the Illumina MiSeq system. The sequences generated with the MiSeq system were performed under accreditation according to the scope of BaseClear B.V. (L457; NEN-EN-ISO/IEC 17025). FASTQ read sequence files were generated using bcl2fastq version 2.20 (Illumina). Initial quality assessment was based on data passing the Illumina Chastity filtering. Subsequently, reads containing PhiX control signal were removed using an inhouse filtering protocol. In addition, reads containing (partial) adapters were clipped (up to a minimum read length of 50 bp). The second quality assessment was based on the remaining reads using the FASTQC quality control tool version 0.11.5.

Paired-end sequence reads were collapsed into so-called pseudoreads using sequence overlap with USEARCH version 9.2 (Edgar, 2010). Classification of these pseudoreads is performed based on the results of alignment with SNAP version 1.0.23 (Zaharia M., 2011) against the RDP database (Cole et al., 2014) version 11.5 for bacterial organisms, while fungal organisms are classified using the UNITE ITS gene database (Abarenkov et al., 2010) version 8.

## Statistical data analysis

Shannon values are calculated with the vegan package using the diversity method. P-values are calculated with the Anova function of the carData package 3.0.2. Results are visualized with ggplot2 version 3.1.1. T-tests with Bonferroni corrected were applied to compare groups. PCA is calculated with the RDA and eigenvals functions of the package vegan. PCA images are visualized with ggplot2 version 3.1.1.

## Results

### Microbiological evaluation

Based on our cytological and bacteriological analysis we could confirm a pyoderma in all four dogs based on *S. pseudintermedius* being present in the skin lesions as pathogen as well as microscopically visible as intracellular bacteria and inflammatory cells within neutrophils.

### Skin Microbiome

Overall, the microbiome of each dog was different but with a substantial overlap (see PCA fig. 1A and B). Between individual dogs the bacterial component showed a larger similarity than the fungal component of the microbiome. In fig. 1C it is observed that the microbiome of the healthy samples was different from the affected samples. The treatment with placebo and the immunoglobulin gel as verum both resulted in a small shift towards the healthy microbiome (not significant possibly due to low number of subjects). There were no specific changes or trends observed in the ITS profiles (Fig. 1D).

**Figure 1:**
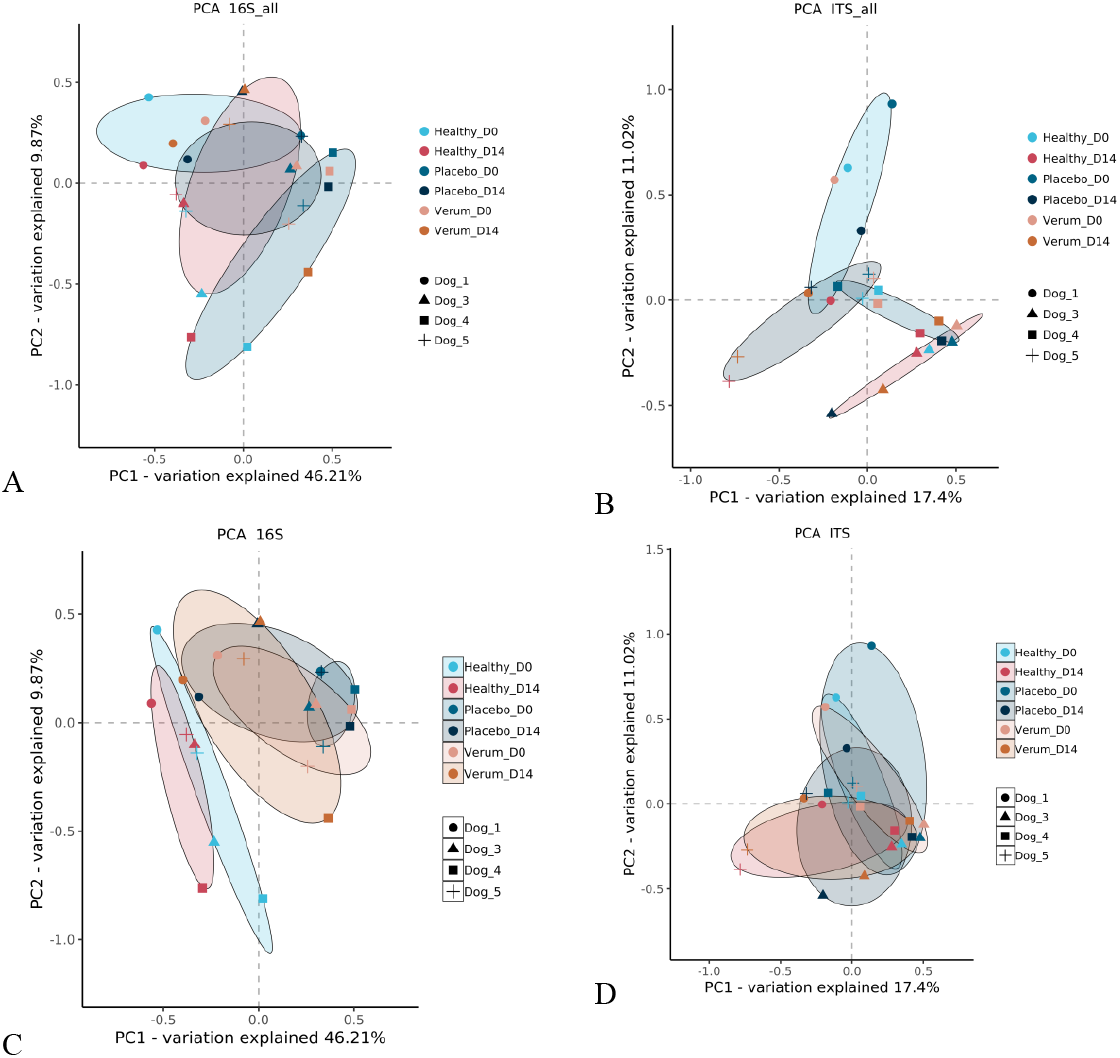
PCA image for the bacterial (16S) and fungal (ITS) component of the microbiome of the dog skin samples. A and B show groups (ellipse) per dog and in C and D the groups indicate timepoint and treatment.

The Shannon diversity (Fig. 2A) was higher in healthy samples, and both verum (4 out of 4 subjects) as well as placebo (2 out of 4 subjects) treatments increased the Shannon diversity of the bacterial component of the microbiome. However, this was not significant (p-values>0.05). For the fungal profiles we did not observe a difference between the sample groups (Fig. 2B) and a similar result was observed in the PCA (Fig. 1D).

**Figure 2A:**
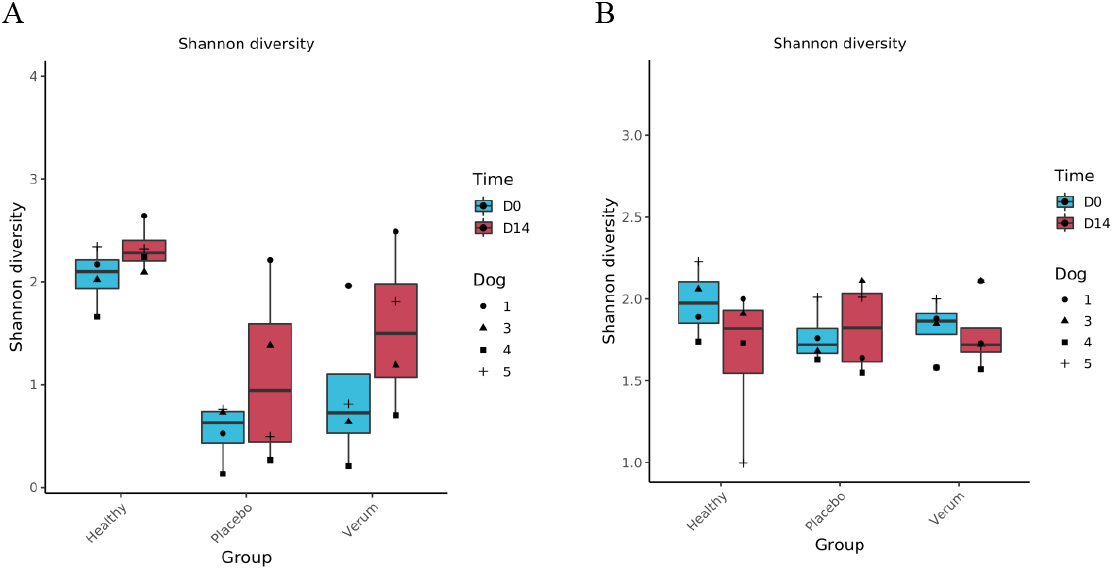
Shannon diversity of the bacterial profiles (based on 16S amplicons), 2B Shannon diversity of the fungal profiles (based on ITS amplicons).

Regarding the relative abundance of *Staphylococcus* there was no significant difference between groups except for day 0 healthy vs placebo p-value=0.016; healthy vs verum day 0 p-value=0.33 which is not significant (see Fig. 3A).

**Figure 3:**
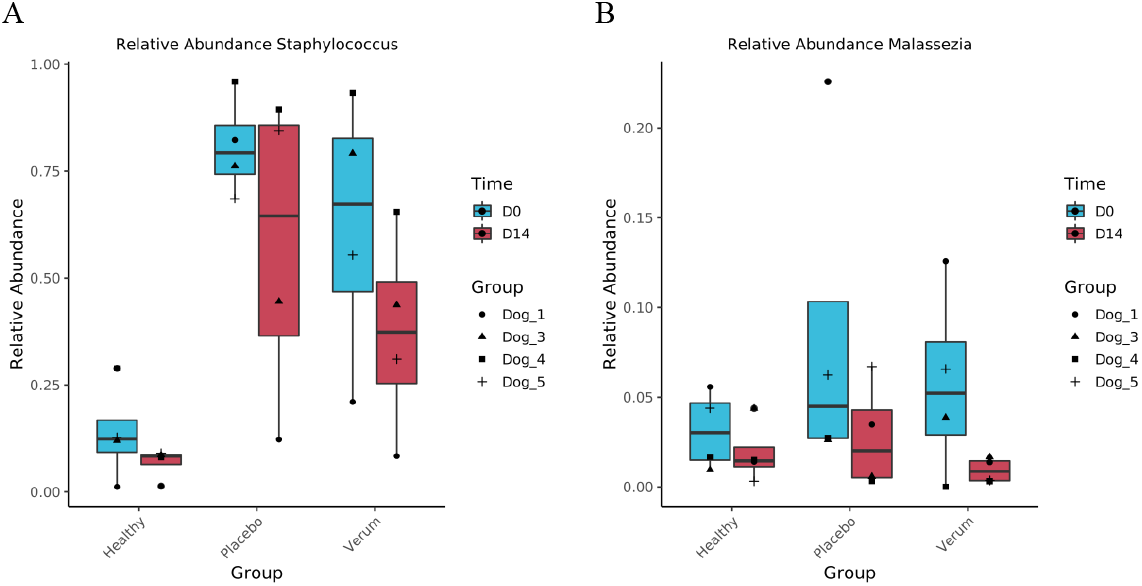
Relative abundance of *Staphylococcus* (A) and *Malassezia* (B) genera in dog skin microbiome samples. Blue: day 0 and Red day 14. •dog 1, Δ dog 3, ▪dog 4, + dog 5.

The % of *Staphylococcus* genera in the skin microbiome of dogs is higher on diseased skin than on healthy skin (see Fig. 3A), however due to the low number of subjects this is not significant. After treatment with verum in four out of four subjects the % of Staphylococcus decreased and this occurred in placebo treatment group in three out of four.

For *Malassezia* no differences are observed between groups. In healthy as well as placebo and verum *Malassezia* slightly decreased in two out of four subjects (no significant findings).

## Discussion

The skin microbiome is a complex ecosystem with important implications for cutaneous health and disease. With our study we could confirm that lesioned canine skin is characterized by a higher percentage of Staphylococci and a lower bacterial diversity in comparison to healthy skin (Chermprapai et al., 2019) and that treatment could shift the microbiome signature of lesioned skin into the direction of healthy skin.

Standard interventions to counteract pathogenic Staphylococci associated with canine pyoderma are antimicrobials such as antibiotics and biocidals in a broad-spectrum manner (Loeffler and Lloyd, 2018). Recently more targeted inventions using phages or phage endolysins have raised interest (Silva et al., 2021). Also, narrow spectrum antimicrobial peptides are discussed as topical alternative for antibiotics (Greco et al., 2019).

Antibodies derived from cow’s milk have an antimicrobial mode of action that is fundamentally different from that of antibiotics and biocides as well as phages and antimicrobial peptides. They do not kill bacteria, but merely disarm them by binding and neutralising their virulence factors. We have shown in previous *in vitro* studies that this leads to a reduction in inflammation and skin damage and stabilises the skin barrier (Geh et al., 2019). In this *in vivo* study, we were able to show that the antibodies have a beneficial influence on the composition of the skin microbiome..

No antibody-dependent specific effect on the proportion of Malassezia in the skin microbiome was measurable. Since Malassezia is both a commensal and a pathogen in cows, we expect antibodies with activity against Malassezia in cow’s milk (Bond, 2010). However, this does not automatically have to result in a reduction in the number of Malassezia but could on the other hand, influence their physiological activity.

The analytical method used in our study does not allow a clear species assignment. This limits the exact assignment of certain species to certain phenomena on the skin as well as effects of the intervention on a particular species. Shotgun metagenome sequencing would allow high coverage for species-level detection as well as functional analysis. Even though the microbial composition can be the same, differences in function and activity can impact the health status. This could be the method of choice for future studies to analyse metabolic differences in the microbiome and the effect of the polyclonal antibody on metabolites.

Both for the bacterial and fungal microbiome, we saw an improvement induced by the gel that acted as carrier of the polyclonal antibody. This is an expected effect of the gel as placebo vehicle that has been described before (Forster et al., 2018).

We consider the demonstrated effects in this pilot study of immunoglobulin from milk in gel on the skin microbiome as promising. However, further studies with an increased number of cases are required to reach statistical significance and zoom into the species and functional levels of the microbiome. We expect to gain more insights into the direct and indirect mode of action of the antibody by binding virulence factors.

## Conclusion

In a pilot study, four dogs diagnosed with generalized superficial pyoderma, received treatment with gels containing a polyclonal antibody isolated from cow’s milk with known activity against virulence factors of *S. pseudintermedius*. The obtained microbiome profiling results indicate that the gel reduced the content of Staphylococcus genera in the skin microbiome and induced an increased overall microbial diversity. Due to the small number of dogs no significant changes were measured. Follow up studies with more subjects are recommended.

## Acknowledgements

The project was funded by the Netherlands antibiotic development platform, grant number NADP20190131LD04.

## Notes

### Competing Interest Statement

The authors have declared no competing interest.

